# Domain Coupling in Allosteric Regulation of SthK Measured Using Time-Resolved Transition Metal Ion FRET

**DOI:** 10.1101/2025.03.31.646362

**Authors:** Pierce Eggan, Sharona E. Gordon, William N. Zagotta

## Abstract

Cyclic nucleotide-binding domain (CNBD) ion channels are vital for cellular signaling and excitability, with activation regulated by cyclic adenosine-or guanosine-monophosphate (cAMP, cGMP) binding. However, the allosteric mechanisms underlying this activation, particularly the energetics that describe conformational changes within individual domains and between domains, remain unclear. The prokaryotic CNBD channel SthK has been a useful model for better understanding these allosteric mechanisms. Here, we applied time-resolved transition metal ion Förster resonance energy transfer (tmFRET) to investigate the conformational dynamics and energetics in the CNBD of SthK in both a soluble C-terminal fragment of the protein, SthK_Cterm_, and in the full-length channel, SthK_Full_. We incorporated the noncanonical amino acid Acd as a FRET donor and a metal bound to a chelator conjugated to a cysteine as an acceptor. We used time correlated single photon counting (TCSPC) to measure time-resolved FRET and fit the TCSPC data to obtain donor-acceptor distance distributions in the absence and presence of cAMP. The distance distributions allowed us to quantify the energetics of coupling between the C-terminal domains and the transmembrane domains by comparing the donor-acceptor distance distributions for SthK_Cterm_ and SthK_Full_. Our data indicate that the presence of the SthK transmembrane domains makes the activating conformational change in the CNBD more favorable. These findings highlight the power of time-resolved tmFRET to uncover the structural and energetic landscapes of allosteric proteins and of the ligand-mediated mechanism in CNBD channels specifically.

## Introduction

Cyclic nucleotide-binding domain (CNBD) ion channels are essential for a variety of physiological processes and are activated when cyclic nucleotides bind to the CNBD of these tetrameric channels (1–3). Upon ligand binding, CNBD channels undergo conformational changes resulting in the opening of the channel pore. However, this pore is over 60 Å away from each of the four peripheral cyclic nucleotide binding sites, and the allosteric mechanisms that underlie this ‘activity at a distance’ to open the pore remain incompletely understood (4–6). The prokaryotic CNBD channel SthK has been a useful model to better understand the underlying mechanisms of ligand activation in these channels. Yet, even with substantial structural and functional information of cyclic adenosine monophosphate (cAMP) binding in SthK (7–11), the mechanism of ligand activation remains unclear from an energetic perspective. As a result, there is a considerable need to integrate both conformational and energetic information in these channels to provide a more complete mechanistic understanding of how cyclic nucleotide binding in the CNBD alters pore opening.

Our previous work utilized time-resolved transition metal ion FRET (tmFRET) to investigate the conformational and energetic changes of the CNBD in an isolated C-terminal fragment of SthK (SthK_Cterm_, Figure 1A, left) (12). tmFRET uses the efficiency of energy transfer between a donor fluorophore and a light absorbing metal ion acceptor to measure the molecular distance between the donor and acceptor. Distance distributions were obtained for the movement of the C-helix relative to the beta-roll of the CNBD in the presence and absence of ligand, which allowed for the calculation of the change in free energy (ΔG) and differences in free energy change (ΔΔG) induced by cyclic nucleotide. However, as these SthK_Cterm_ measurements were made in a truncated fragment of the protein, it remained unclear how the CNBD conformational change might look in the full-length channel (SthK_Full_, Figure 1A, right) and how the energetics of this transition might differ between the full channel and SthK_Cterm_.

**Figure 1.**
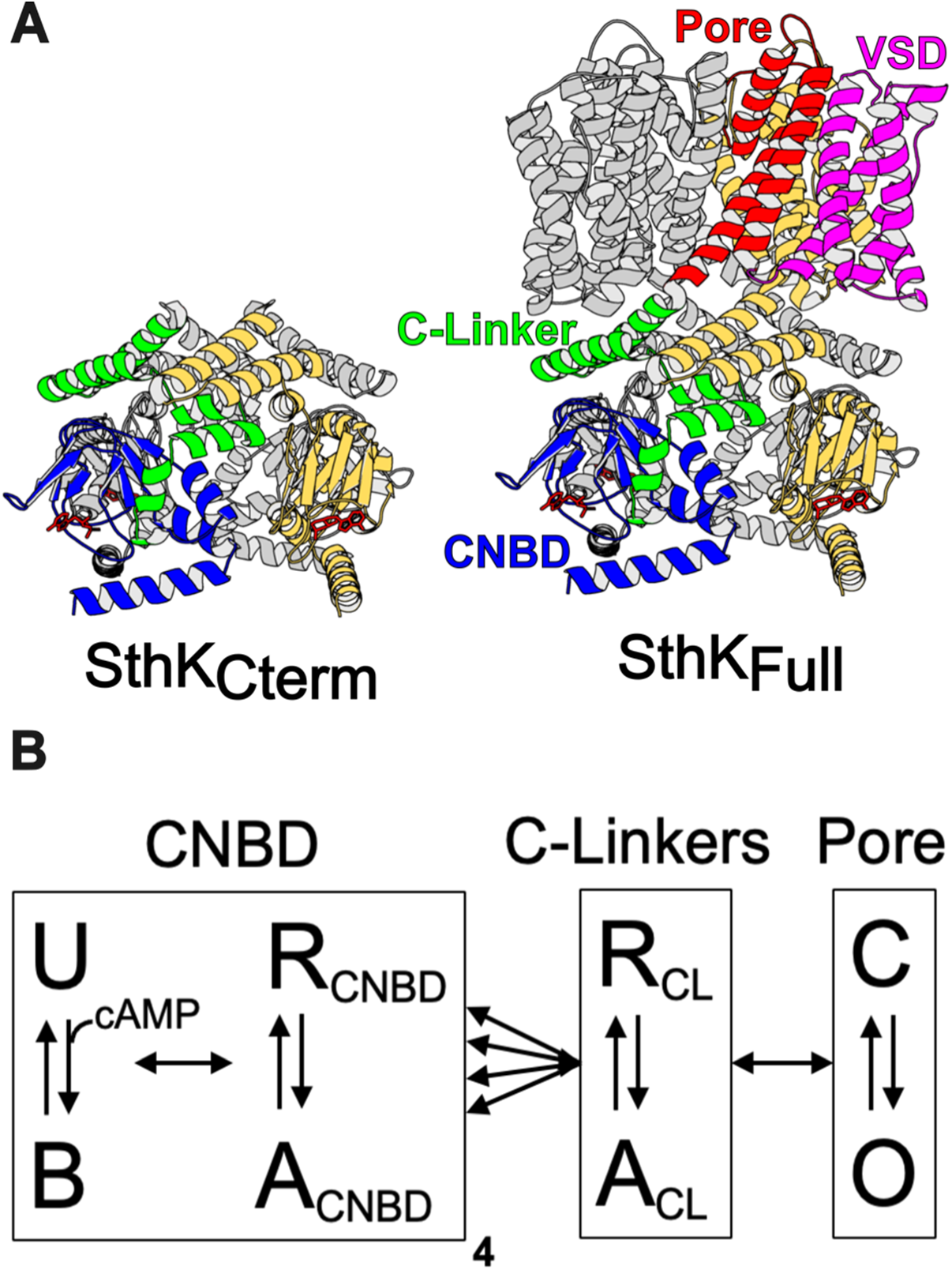
Structure of SthK constructs and modular gating scheme. **(A)** Cartoon structures for SthK_Cterm_ and SthK_Full_ with the domains labeled on the SthK_Full_ structure. **(B)** Modular gating scheme for SthK showing modules corresponding to domains indicated in the structures above. Within each domain, resting (R) and active (A) or closed (C) and open (O) or unbound (U) and bound (B) conformations are shown connected by double arrows, representing the transition between states. Horizontal arrows between modules indicate the coupling between domains.

We approach this question using a modular gating scheme, also known as an ensemble allosteric model, which provides a framework for integrating structural and energetic contributions across protein regions, as illustrated in Figure 1B (13–15). In this model, allosteric transitions occur through coupled conformational changes in defined modules, which for SthK and related CNBD channels, we define as the structural domains of the pore, C-linker, and CNBDs (Figure 1A). We depict four independent CNBD transitions and concerted transitions in the C-linker and in the pore, although alternative arrangements and connections are possible (5, 16, 17). Each module has a conformational equilibrium between its resting state (R) and active state (A) (vertical arrows in Figure 1B) that is described by a corresponding free energy change, ΔG. Importantly, these transitions in individual modules are not independent but are coupled to each other (horizontal arrows in Figure 1B). Coupling is the amount by which the energetics of a module’s transition is changed by the state of the neighboring module (ΔΔG). Ultimately, understanding the energetics of all of the arrows in Figure 1B is essential to understand the mechanisms of CNBD channel gating.

Quantifying the ΔGs and ΔΔGs for SthK_Cterm_ and SthK_Full_ requires a technique that can measure the conformational changes and energetics of each individual module. Time-resolved tmFRET offers insights into protein conformational and energetic changes by using fluorescence lifetimes to quantify distance distributions between a donor fluorophore and a metal ion acceptor. Importantly, our tmFRET method is applicable to both soluble and membrane protein structures under physiological conditions, making it ideal for studying SthK dynamics and membrane proteins more broadly. The theoretical framework of time-resolved tmFRET has been previously described (12, 18–21).

To apply the time-resolved tmFRET approach to full-length SthK channels, we first extended our method to measuring lifetimes in the time-domain using time-correlated single photon counting (TCSPC). We believe that TCSPC provides several advantages over our previously utilized frequency-domain measurements on a fluorescence lifetime imaging microscope (FLIM). As membrane protein expression and purification is challenging, we needed an approach that allowed us to use much lower concentrations of protein than required for our previous experiments. The TCSPC cuvette-based instrument has a longer pathlength than on an imaging microscope and therefore allowed us to use ∼10-fold lower protein concentrations.

Additionally, unlike frequency domain measurements, the noise in TCSPC data is due to single-photon shot noise and thus can be well described by a Poisson distribution (above 10 photons), making the χ^2^ determinations of model fits more meaningful (21). Lastly, TCSPC allows for better accounting of the background fluorescence by direct measurement, which allows for more reliably fitting data regardless of protein concentration or buffer conditions.

As in the frequency domain experiments, lifetimes measured with TCSPC decrease in the presence of FRET. These data can be fit by a model with a distribution of distances that describes the measured FRET between donor and acceptor molecules in the sample. This distance distribution accounts for heterogeneity within and between conformational states of the protein, the latter of which is directly related to the energetics of conformational transitions through the Boltzmann relationship (12, 21).

In this study, we used time-resolved tmFRET with TCSPC data to measure the ligand-induced conformational changes in the CNBD of SthK, first in the isolated C-terminal fragment, SthK_Cterm_, and then in the full-length channel, SthK_Full_. Distance distributions were obtained for both SthK_Cterm_ and SthK_Full_ in the presence and absence of ligand. This allowed for calculation of

ΔG_apo_ and ΔG_cAMP_ in each construct and the determination of the energetic contribution of coupling to the pore domain. These findings additionally validate the use of TCSPC data in time-resolved tmFRET and demonstrate its capability to measure unique distance distributions across different protein constructs for energetic analysis. Ultimately, these results contribute to a deeper understanding of the allosteric mechanism of ligand gating in SthK.

## Results

### Validation of Time-Resolved tmFRET Using TCSPC in SthK_Cterm_

To first validate our time-resolved tmFRET approach using TCSPC, we started with our previously published construct of the C-terminal fragment of SthK (SthK_Cterm_) (12). This allowed us to compare distance distributions obtained with TCSPC with those previously obtained with frequency domain fluorescence lifetime imaging microscopy (FLIM) in the same protein construct. In these experiments, SthK_Cterm_, comprised of the isolated C-linker domain and CNBD, was expressed with the noncanonical amino acid acridon-2-ylalanine, Acd, at S361 and a cysteine at V416 for metal-ion acceptor conjugation. As previously described, SthK_Cterm_ incorporating Acd was purified and mixed with excess WT unlabeled SthK_Cterm_, to create heterotetrameric SthK_Cterm_:WT/S361Acd-V416C protein (cartoon in Figure 2A). This heteromeric channel eliminated the possibility of FRET between neighboring subunits (12). In parallel, SthK_Cterm_:WT/S361Acd-V416C (with a cysteine) and SthK_Cterm_:WT/S361Acd (donor-only, without any cysteines) protein constructs were incubated with the thiol-specific metal ion acceptor [Ru(bpy)_2_phenM]^2+^. This acceptor has an *R*_O_ = 43.5 Å when paired with Acd, where *R*_O_ is the distance at which FRET efficiency is 0.5 (structures of donor and acceptor labels shown in Figure 2A) (22). The labeled protein was subjected to size exclusion chromatography (SEC) to remove unreacted [Ru(bpy)_2_phenM]^2+^ as well as any monomeric protein, and the purified tetrameric protein was used immediately in fluorescence lifetime experiments.

**Figure 2.**
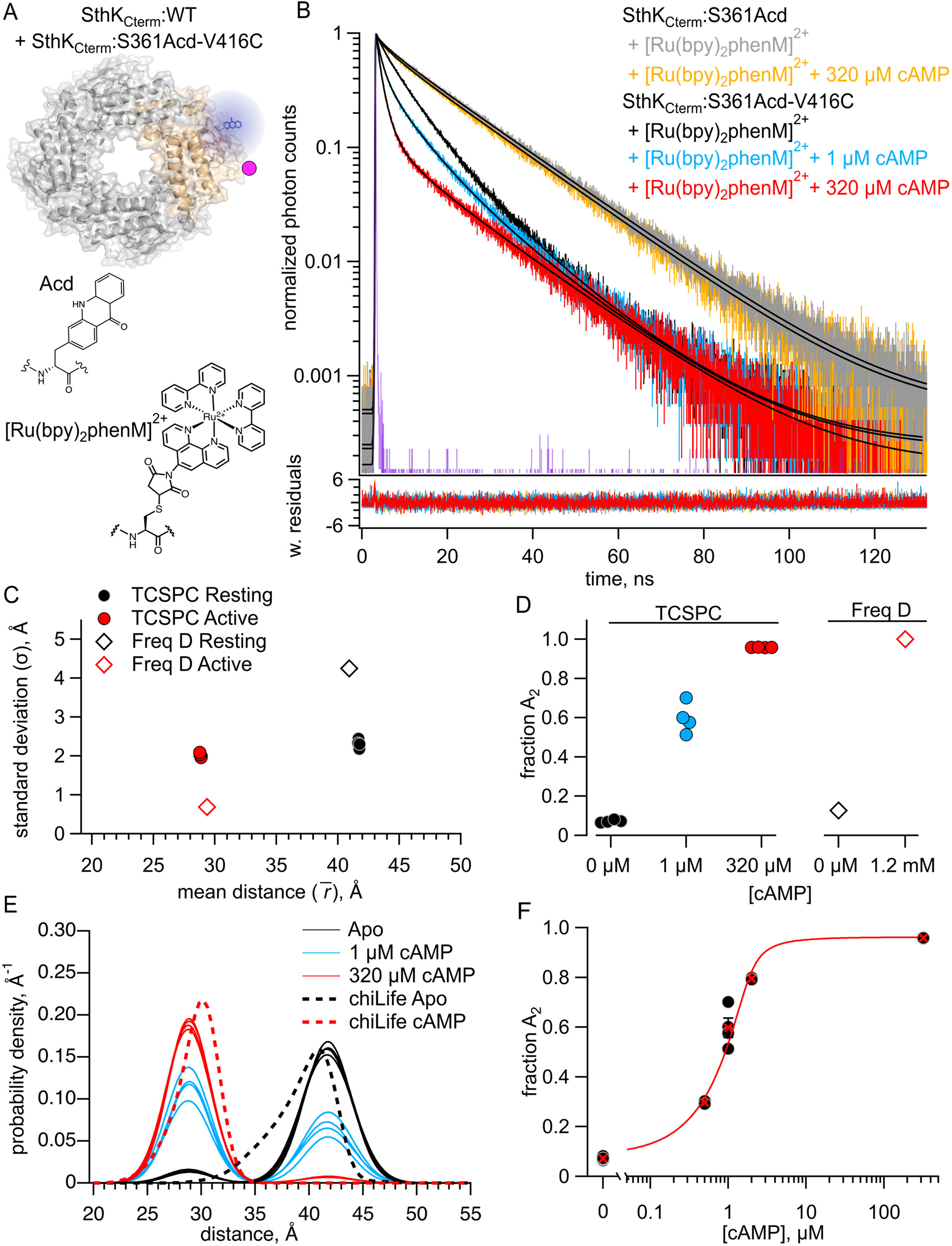
FRET with the donor-acceptor pair within the same subunit. (**A**) Cartoon of SthK_Cterm_:WT/S361Acd-V416C with a single donor-acceptor labeled subunit (top; acceptor shown as magenta dot) and the structures of donor fluorophore, Acd, and acceptor metal complex [Ru(bpy)_2_phenM]^2+^ (bottom). (**B**) Representative TCSPC data from SthK_Cterm_:WT/S361Acd treated with [Ru(bpy)_2_phenM]^2+^ before (grey) and after (orange) the addition of 320 μM cAMP. TCSPC data from SthK_Cterm_:WT/S361Acd-V416C treated with [Ru(bpy)_2_phenM]^2+^ in the absence of cAMP (black) and in the presence of 1 μM (blue) or 320 μM (red) cAMP. Smooth black curves show the fits of the FRET model, and weighted residuals are shown below using the same colors as used for the data. Representative instrument response function (IRF) shown in purple. (**C**) Summary of distance parameter values, 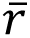 and σ (n = 4), from fits to our TCSPC data (filled symbols) and from previously published frequency domain data (open symbols), with resting and active state as indicated in the legend (12). (**D**) *f*_A2_ values determine from fits to TCSPC data measured with 0, 1 μM and 320 μM cAMP (filled symbols) compared to previously published *f*_A2_ values determine from frequency domain data (open symbols) (12). (**E**) Spaghetti plot of distance distributions from individual experiments for 0 (black), 1 μM (blue) and 230 μM (red) cAMP, with distributions predicted by chiLife shown as dashed curves. Resting and active are shown in black and red, respectively. (**F**) Summary of *f*_A2_ values over range of cAMP concentrations. The dose-response relation was fit with a quadratic equation (red curve) with *K_D_* = 0.22 μM and [*P*]_total_ = 1.2 μM.

We collected fluorescence lifetime data in the form of TCSPC decays. First, the donor-only SthK_Cterm_:WT/S361Acd was measured and the photon arrival times are shown on a normalized log scale histogram (grey trace, Figure 2B). We observed that this lifetime, while approximating a single-exponential decay, was best fit with a double exponential decay, with 87% of the amplitude arising from a 17.6 ns lifetime and the remaining contribution from a second 4.73 ns lifetime. The SthK_Cterm_:S361Acd donor-only protein lifetime did not significantly change in the presence of cAMP, as expected (orange trace, Figure 2B). This lifetime is very similar to that previously described for Acd in the same protein using frequency domain lifetime measurements (12). These donor-only lifetimes for each condition were then used as fixed parameters when analyzing the amount of FRET in our cysteine-containing protein.

We next measured lifetimes of the SthK_Cterm_:WT/S361Acd-V416C-[Ru(bpy)_2_phenM]^2+^ protein, in the absence of ligand (black trace, Figure 2B). As expected, labeling this construct with [Ru(bpy)_2_phenM]^2+^ decreased the overall lifetime of Acd compared to the donor-only lifetime because of FRET. In addition to decreasing, the lifetime of acceptor-labelled protein became even more nonexponential, indicative of multiple donor-acceptor distances contributing to the FRET. Subsequently, upon the addition of a subsaturating concentration of 1 µM cAMP, the lifetime of Acd decreased (blue trace, Figure 2B) and upon addition of a saturating concentration of 320 µM cAMP decreased yet further (red trace, Figure 2B). This decrease in the fluorescence lifetime indicates increased FRET and shorter distances between the donor and acceptor molecules in response to cAMP. Overall, these cAMP-induced decreases in lifetimes, compared with our control without acceptor, demonstrate that our TCSPC approach has the sensitivity to measure ligand-triggered conformational changes in the CNBD of SthK_Cterm_.

We fit the SthK_Cterm_:WT/S361Acd-V416C-[Ru(bpy)_2_phenM]^2+^ lifetimes with a model that assumes a distribution of donor-acceptor distances. We parameterized the distance distribution as the sum of two Gaussians, each with an average distance (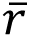) and a standard deviation (σ), as well as fraction *A_2_* (*f*_A2_) describing the relative contribution of each Gaussian. These two Gaussians represent the resting and active conformational states of the CNBD C-helix. This FRET model was the same as was used for our previously published frequency domain measurements (12, 19–21), but adapted to TCSPC lifetime measurements (see Materials and methods.)

We globally fit the lifetimes from apo and a range of cAMP concentrations (0.5, 1, 2, 320 µM; Figure 2B). These fits converged with a reduced χ^2^ near 1, and the parameters were well-identified (see Materials and methods). The 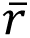 and σ values for the resting and active state Gaussians from different experiments are summarized in Figure 2C, with an average distance 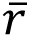_1_ = 41.7 Å and σ_1_ = 2.3 Å in the resting state and 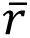_2_ = 28.8 Å and σ_2_ = 2.1 Å in the active state. These values were similar to our previously published frequency domain results of this same donor-acceptor, however the σ values showed less state dependence (Figure 2C) (12). The fraction of the distribution in the active state, *f*_A2_, was best fit with 8% in the absence of ligand and 96% in the presence of a saturating concentration of cAMP, again similar to our previous frequency domain measurements (Figure 2D). A summary of distributions from individual experiments across various cAMP concentrations is shown in the spaghetti plot in Figure 2E. Overlayed on the spaghetti plots are the distance distributions predicted by the known structures using chiLife software (12, 23). These computationally predicted distance distributions show a remarkable similarity to those based on fits to our experimental data. The *f*_A2_ values for a range of cAMP concentrations are displayed as a dose-response curve, which was fit with a quadratic equation assuming no binding cooperativity (K_D_ = 0.22 µM; Figure 2F). This K_D_ value is comparable to the value obtained previously using steady-state tmFRET (12). Based on these data, TCSPC determined distance distributions at least as well as our previous frequency domain approach.

### Quantifying Intersubunit FRET in SthK_Cterm_

Our ultimate goal of examining conformational energetics in full-length SthK, which includes the membrane spanning domains, presents an additional experimental challenge. Unlike in the C-terminal fragment in which it is straightforward to make heterotetramers including only a single labeled subunit, at this time we can only express and purify full-length SthK as homotetramers.

These homotetramers would, of necessity, have donors and acceptors in all subunits and thus could have intersubunit FRET. To address the contribution of intersubunit FRET, we measured lifetimes in a heterotetramer with Acd incorporated into only a single SthK_Cterm_:S361Acd subunit (with no acceptor cysteine) and [Ru(bpy)_2_phenM]^2+^ acceptors on the remaining three SthK_Cterm_:V416C subunits (with no Acd donor fluorophores). This protein was produced by mixing an excess of SthK_Cterm_:V416C protein with SthK_Cterm_:S361Acd (Figure 3A). We measured shorter lifetimes in these heterotetramers than observed for control donor-only protein, both in the absence and presence of cAMP (Figure 3B, black and red traces, respectively). Fits to these data with the FRET model gave parameter values of 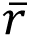_1_= 54.1 Å and σ_1_ = 3.0 Å in the resting state, and 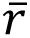_1_ = 58.0 Å and σ_1_= 5.0 Å in the active state (Figure 3C). The resting and active distributions aligned well with the intersubunit distance distributions predicted by chiLife (Figure 3D). Indeed, the fits most closely aligned with the shortest predicted intersubunit distance, conforming to expectations that the closest distances would dominate the FRET. Based on these results, we conclude that intersubunit FRET must be considered when fitting data from experiments with homotetrameric SthK with all four subunits labeled with donor and acceptor.

**Figure 3.**
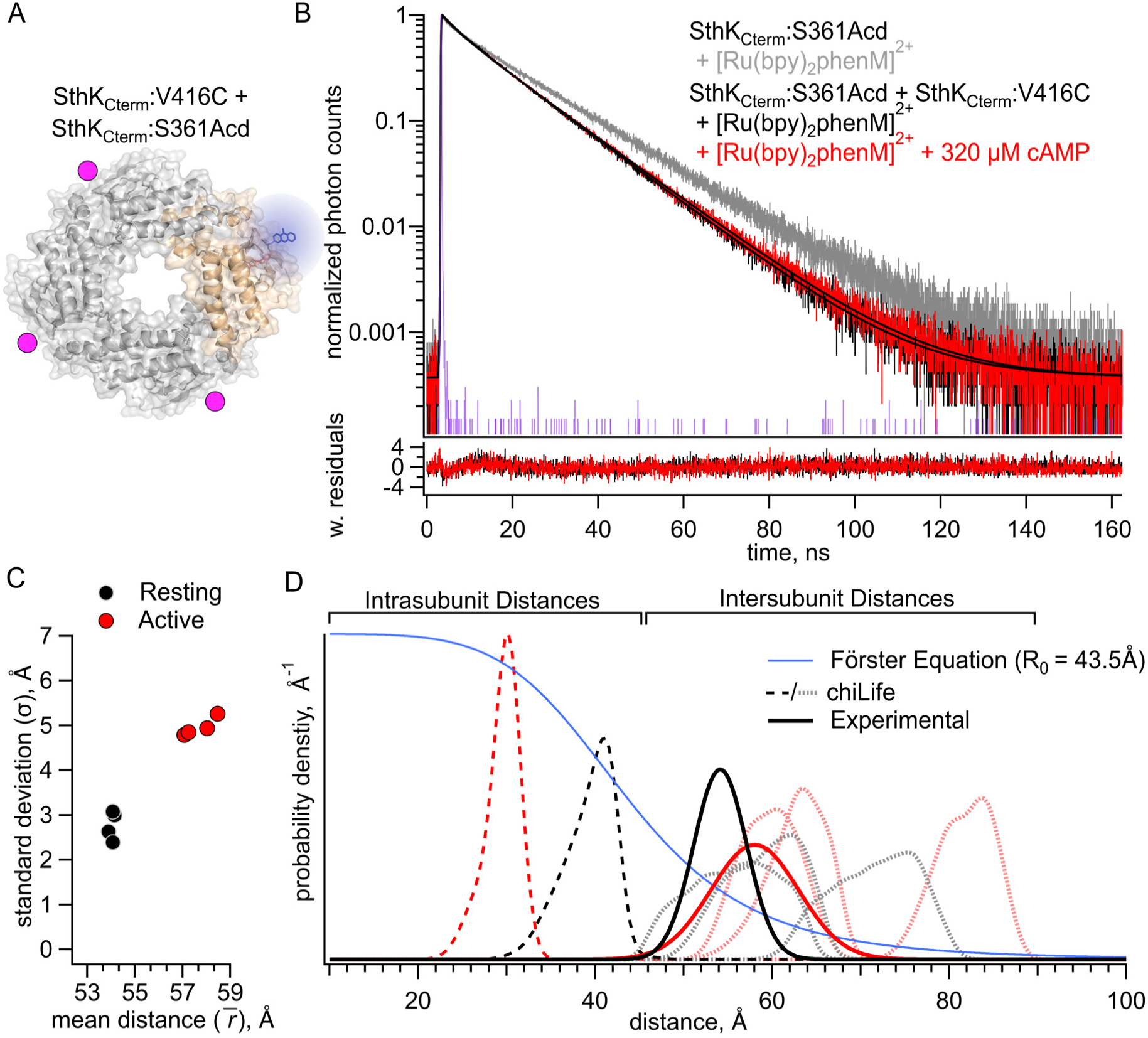
FRET with the donor-acceptor pair on different subunits. **(A)** Cartoon structure of SthK_Cterm_:V416C/ S361Acd protein with a single donor-labeled subunit (SthK_Cterm_:S361Acd) and 3 acceptor-labeled subunits (SthK_Cterm_:V416C). **(B)** Representative TCSPC data from the donor-only condition (grey), and after [Ru(bpy)_2_phenM]^2+^ labeling either in the absence (black) or presence (red) of 320 μM cAMP, fit with the FRET model (smooth black lines). The IRF and weighted residuals are depicted as in Figure 2. **(C)** Summary of distance parameter values, 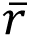 and σ (n = 4), from fits of our TCSPC data with the FRET model. Resting and active state are shown as black and red, respectively. **(D)** Distributions of donor-acceptor distances predicted with chiLife for intrasubunit distances (darker dashed curves) and intersubunit distances (lighter dashed curves), with black and red as described in (C). The distance distributions from fits to our TCSPC data for resting and active states are overlayed as solid black and red Gaussians.

### Measuring homotetrameric SthK_Cterm_ with both intersubunit and intrasubunit FRET

After quantifying the intersubunit FRET in heterotetramers that had FRET only between subunits, we proceeded to measure fluorescence lifetimes in a homotetrameric construct that contained both intrasubunit and intersubunit FRET (Figure 4A). We measured lifetimes for homomeric SthK_Cterm_:361Acd-V416C-[Ru(bpy)_2_phenM]^2+^ in the absence of cAMP and in the presence of either 1 or 320 µM cAMP (Figure 4B, black, blue, and red traces, respectively). To account for the contributions from intersubunit FRET, our analysis added a second FRET acceptor with the experimentally determined intersubunit distance distribution (Figure 3C, see Materials and methods). Global fits of this FRET model to the data gave intrasubunit distances of 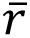_1_ = 41.6 Å (σ_1_= 3.6 Å) and 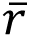_2_ = 29.1 Å (σ_2_= 4.1 Å), for the resting and active states, respectively (Figures 4B and 4C). These 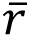 values were similar to those obtained from heteromeric SthK_Cterm_:WT/S361Acd-V416C-[Ru(bpy)_2_phenM]^2+^, with somewhat wider σ values (see Figure 2C). Interestingly, the distribution among states in the homotetramer, with a *f*_A2_ = 0.25 in the absence of ligand, and *f*_A2_ = 0.94 in the presence of a saturating concentration of ligand, was somewhat different from the SthK_Cterm_:WT/S361Acd-V416C-[Ru(bpy)_2_phenM]^2+^ heterotetramers (Figure 4D vs. Figure 2D). This suggests a slightly more favorable closing of the C-helix in the absence of cAMP and slightly less favorable closing in the presence of saturating cAMP in homotetramers. A summary of distance distributions from different experiments across cAMP concentrations is shown as a spaghetti plot in Figure 4E, with the chiLife predictions overlayed as dashed curves. Based on these data, we conclude that there are only minimal differences in the distribution among resting and active states between the homotetrameric and heterotetrameric protein.

**Figure 4.**
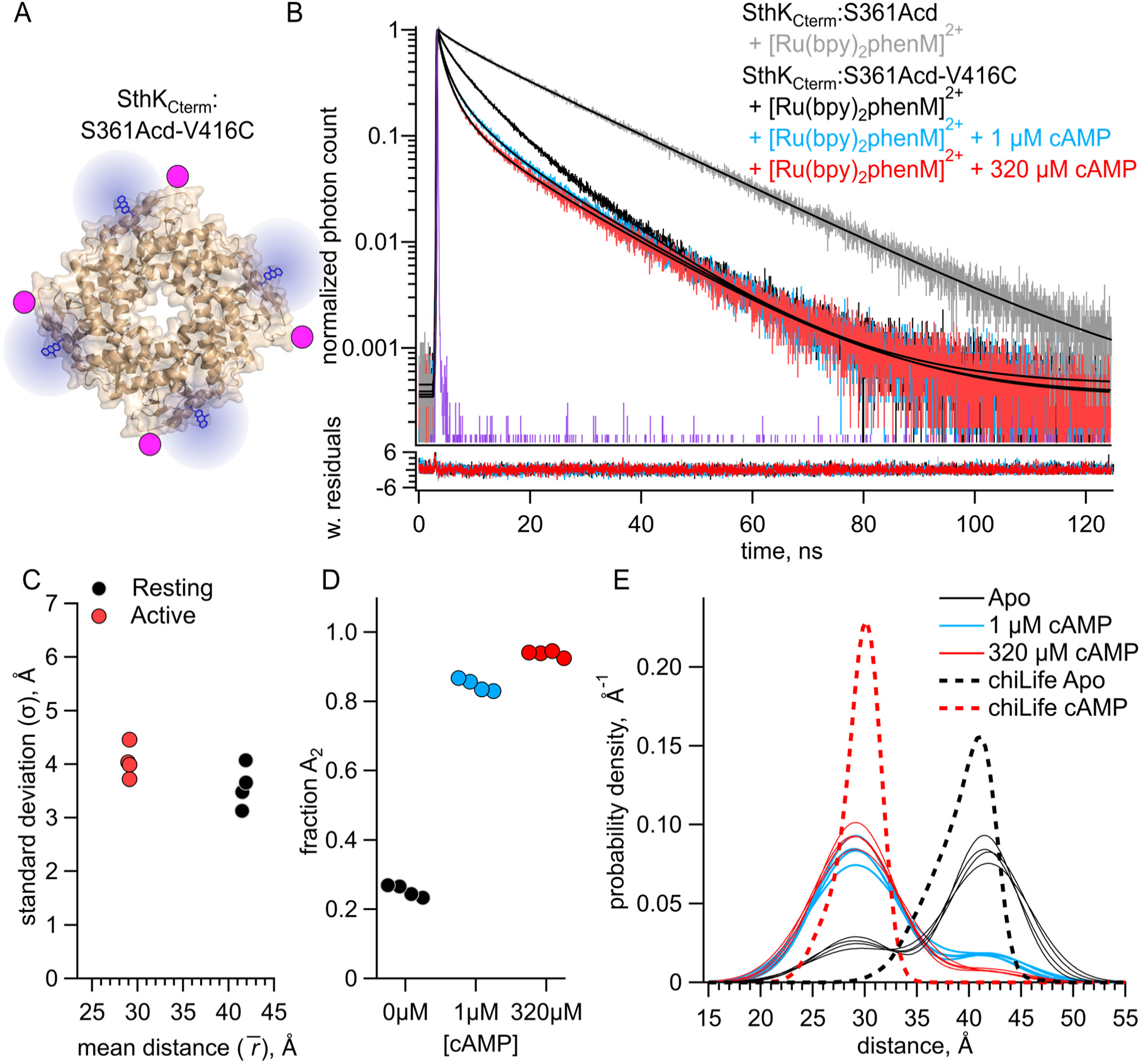
FRET with both intrasubunit and intersubunit donor-acceptor pairs in SthK_Cterm_. **(A)** Cartoon structure of homotetrameric SthK_Cterm_:S361Acd-V416C protein. **(B)** Representative TCSPC data from the donor-only condition (in grey) and after [Ru(Bpy)_2_phenM]^2+^ labeling in the absence of cAMP (black) and in the presence of either 1 μM (blue) or 320 μM (red) cAMP, fit with the FRET model (smooth black lines). The IRF and weighted residuals are depicted as in Figure 2. **(C)** Summary of distance parameter values, 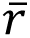 and σ (n = 4), from fits of our TCSPC data with the FRET model. Resting and active state are shown as black and red, respectively. **(D)** Values of *f*_A2_for 0, 1 μM or 320 μM cAMP, as indicated. **(E)** Spaghetti plot of distance distributions from individual experiments for 0 (black), 1 μM (blue) and 230 μM (red) cAMP, with distributions predicted by chiLife shown as dashed curves. Resting and active are indicated by black and red, respectively.

### Full-Length SthK Acd Incorporation and Purification

Having established a method to account for intersubunit FRET, we expressed and purified full-length SthK protein (SthK_Full_) using the same Acd site at residue 361, either with or without a cysteine at the same acceptor position 416. These constructs were expressed in *E.coli*, where membrane fractions were isolated and solubilized in n-Dodecyl-β-D-maltoside (DDM) detergent with 10:1 cholesteryl hemisuccinate (CHS). The SthK_Full_ was then purified using Strep-Tactin affinity resin, and the DDM detergent was exchanged for Lauryl Maltose Neopentyl Glycol (LMNG) detergent + CHS on the Strep-Tactin column (see Materials and methods).

SthK_Full_:S361Acd-V416C (with cysteine) and SthK_Full_:S361Acd (without cysteine) were successfully purified as confirmed with in-gel fluorescence (Figure 5A). Coomassie staining showed a small proportion of protein subunits (< 15%) that appeared truncated at the TAG codon site and copurified with full-length subunits. Size exclusion chromatography showed predominantly a single peak in Acd fluorescence without aggregates, indicating monodispersed well-behaved protein (Figure 5B).

**Figure 5.**
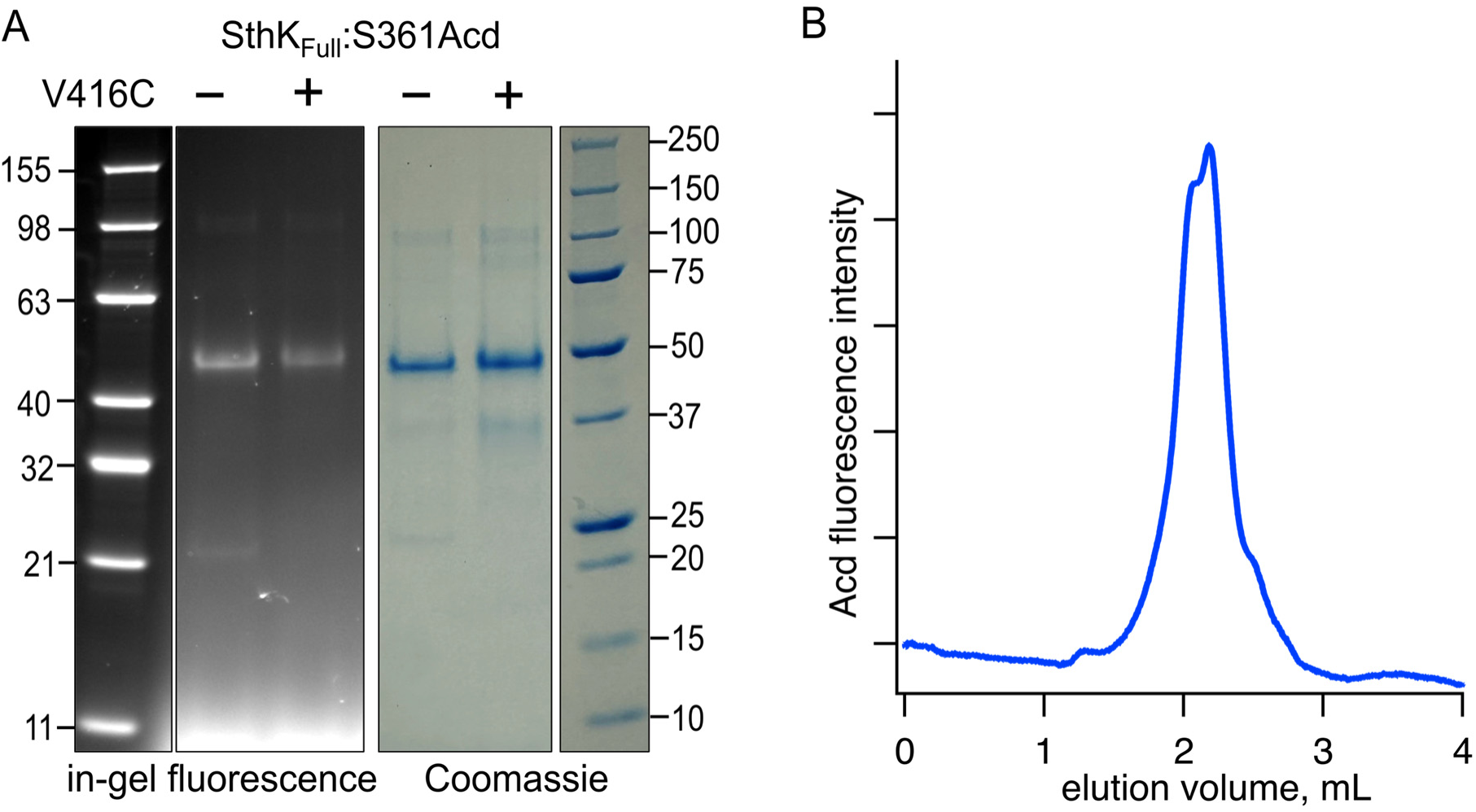
Expression and purification of SthK_Full_ incorporating Acd. **(A)** SDS/PAGE with in-gel fluorescence (left) or Coomassie blue (right) for SthK_Cterm_:S361Acd and SthK_Cterm_:S361Acd-V416C, as indicated (theoretical mass = 53.0 kDa). **(B)** Chromatogram from size exclusion chromatography monitoring fluorescence at 425 nm (Acd) for SthK_Cterm_:S361Acd-V416C.

### Measuring tmFRET in Full-Length SthK

We used TCSPC to measure the fluorescence lifetimes of both purified constructs, SthK_Full_:S361Acd-V416C and SthK_Full_:S361Acd, before and after incubating with [Ru(bpy)_2_phenM]^2+^ acceptor (Figure 6A). The lifetime of SthK_Full_:S361Acd donor-only protein was similar to that of SthK_Cterm_:S361Acd (Figure 6B, gray trace), and the addition of cAMP did not change the lifetime (Figure 6B, orange trace). As expected, the lifetime of SthK_Full_:S361Acd-V416C became shorter upon labeling with [Ru(bpy)_2_phenM]^2+^ (Figure 6B, black). The addition of a range of cAMP concentrations (0.25, 0.5, 1, 2 and 320 µM) to SthK_Full_:S361Acd-V416C-[Ru(bpy)_2_phenM]^2+^ protein further decreased the lifetimes, following the same trend as was observed in the SthK_Cterm_ protein (Figure 6B).

**Figure 6.**
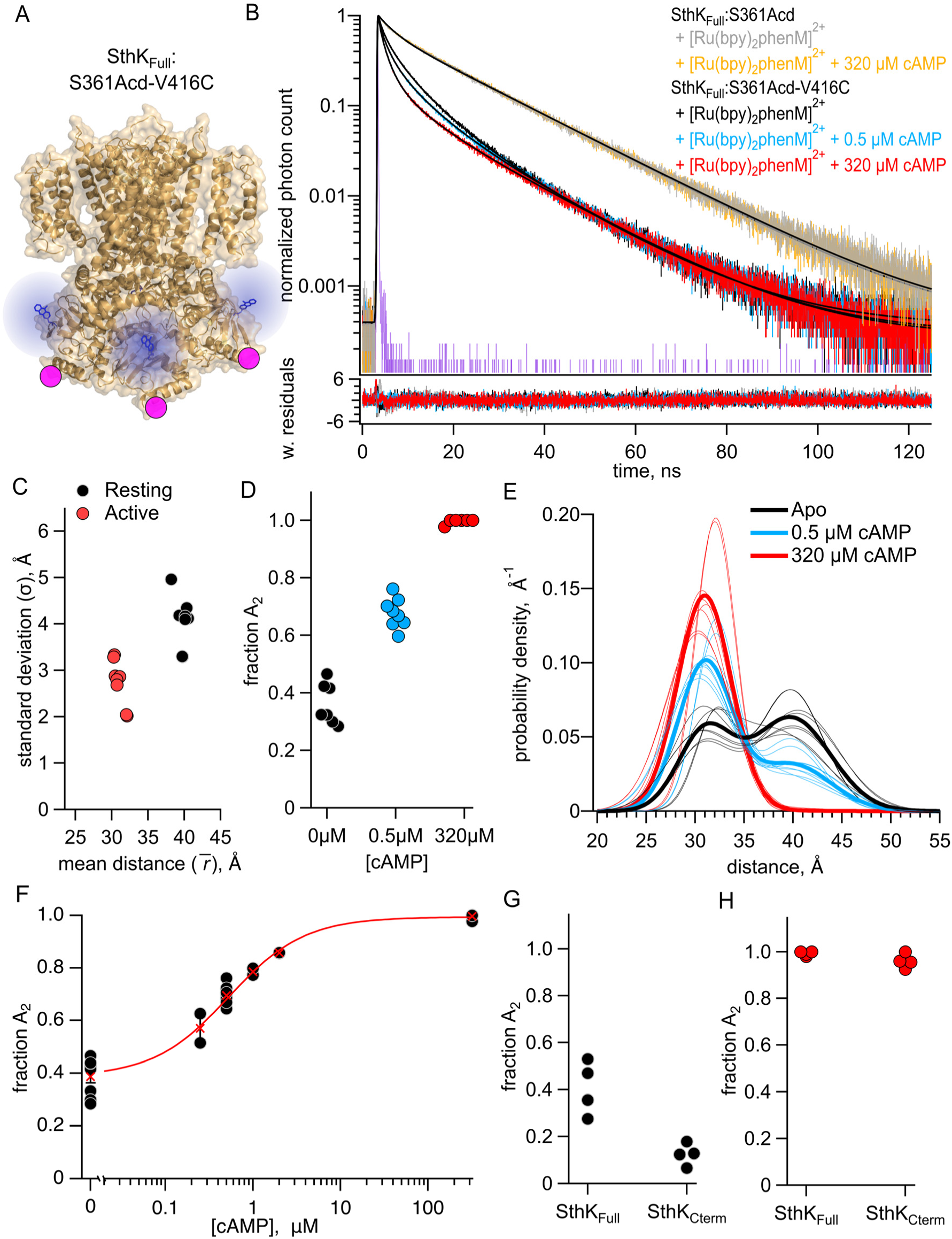
FRET with both intrasubunit and intersubunit donor-acceptor pairs in SthK_Full_. **(A)** Cartoon structure of SthK_Full_:S361Acd-V416C construct. **(B)** Representative TCSPC data from the donor-only condition (no cAMP – gray, 320 µM cAMP – orange) and SthK_Full_:S361Acd-V416C after [Ru(bpy)_2_phenM]^2+^ labeling in the absence of cAMP (black) and in the presence of either 1 μM (blue) or 320 μM (red) cAMP, fit with the FRET model (smooth black lines). The IRF and weighted residuals are depicted as in Figure 2. **(C)** Summary of distance parameter values, 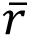 and σ (n = 4), from fits of our TCSPC data with the FRET model. Resting and active state are shown as black and red, respectively. **(D)** Values of *f*_A2_ for 0, 1 μM or 320 μM cAMP, as indicated. **(E)** Spaghetti plot of distance distributions from individual experiments for 0 (black), 1 μM (blue) and 230 μM (red) cAMP, with distributions predicted by chiLife shown as dashed curves. Resting and active are indicated by black and red, respectively. **(F)** Summary of *f*_A2_ over range of cAMP concentrations. The dose-response relation was fit with a quadratic equation (red curve) with *K_D_* = 0.53 μM and [*P*]_total_ = 0.025μM. **(G-H)** Values of *f*_A2_ from globally fitting SthK_Full_ and SthK_Cterm_ constructs.

We globally fit the lifetime data from SthK_Full_ with the FRET model that includes the intrasubunit and intersubunit FRET contributions (Figure 6B). We found that the best fit for the dataset had average parameter values of 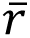_1_ = 39.8 Å (σ_1_= 4.1 Å) and 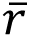_2_ = 31.0 Å (σ_2_ = 2.7 Å) for the resting and active state Gaussians, respectively (Figure 6C). These parameter values are similar to those obtained from homotetrameric SthK_Cterm_, particularly for 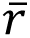s, which were within 1.5 Å of previous values (Figure 4C). The *f*_A2_ values, however, showed SthK_Full_ had a higher probability of being in the active state, both in the absence of cAMP and in the presence of a saturating concentration of cAMP, compared to SthK_Cterm_ (Figure 6D). The distance distributions for the different SthK_Full_ experiments are summarized in spaghetti plots in Figure 6E. Across a range of subsaturating cAMP concentrations, *f*_A2_showed intermediate values between those obtained from apo and saturating cAMP (Figure 6F). This relationship was fit with a quadratic equation to give a K_D_ = 0.52 µM, which is comparable to that obtained from the SthK_Cterm_ but substantially lower than the K_D_ reported from WT-SthK electrophysiology experiments (K_D_ = 1.5 µM) (24). These values, however, should be interpreted with caution due to the CNBD’s high affinity for cAMP and the challenges of accurately determining the total protein concentrations in our experiments. Overall, in SthK_Full_ containing the pore and transmembrane domains and in a detergent environment, we observed similar resting and active state distances (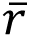 and σ) for the CNBD C-helix, but different relative proportions between these states (*f*_A2_) compared to the SthK_Cterm_.

To focus on changes in *f*_A2_in SthK_Full_ compared to SthK_Cterm_, we globally fit the SthK_Full_ and SthK_Cterm_ data assuming the resting and active states average distances (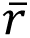) and standard deviations (σ) were the same across both proteins, and only *f*_A2_was allowed to vary. We believe this assumption is reasonable based on the similar distance distributions we found for SthK_Full_ and SthK_Cterm_ (Figure 6C and 2C) and the similar structures of the full-length SthK channel in the cAMP bound and active state (PDB: 7RTJ) and the C-terminal SthK with cAMP bound (PDB:4D7T) (25, 26). Our global fit included data in the absence of cAMP and a saturating concentration of cAMP for both SthK_Full_ and SthK_Cterm_ and used the FRET model with both intersubunit and intrasubunit FRET. The best fit to both datasets yielded 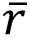_1_ = 40.5 Å with σ_1_= 4.1 Å and 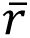_2_ = 30.5 Å with σ_2_= 3.3 for the absence of cAMP and a saturating concentration of cAMP, respectively. As observed for independent fits to each dataset (Figure 4D and Figure 6D), the globally fit *f*_A2_differed between the protein constructs, with SthK_Full_ (apo *f*_A2_= 0.41, cAMP *f*_A2_ = 0.99) giving a higher percentage of active conformations in both the absence of cAMP and the presence of a saturating concentration of cAMP, compared to SthK_Cterm_ (apo *f*_A2_ = 0.12, cAMP *f*_A2_= 0.96) (Figure 6G and H).

The proportion of molecules in each state can be converted into the change in Gibbs free energy (ΔG) for the CNBD transition in the modular gating scheme in Figure 1B. A summary of the energetics for this conformational transition of the C-helix, based on the global fits across the SthK_Cterm_ and SthK_Full_ datasets, is shown in Figure 7. In the SthK_Cterm_ protein construct, the resting to active conformational transition is unfavorable in the absence of ligand (ΔG = +1.15 kcal/mol) and is more favorable in the presence of saturating cAMP (ΔG =-1.87 kcal/mol). In comparison, in SthK_Full_ with attached pore and transmembrane domains, the resting to active transition without ligand is still unfavorable (ΔG = + 0.22 kcal/mol), but more favorable than in SthK_Cterm_, and cAMP further stabilizes the transition (ΔG =-2.87 kcal/mol). This suggests coupling between the pore and the cytoplasmic domain stabilizes the active state of the CNBD by about-1 kcal/mol. According to the modular gating scheme, domain coupling should not affect the difference in change of free energy (ΔΔG) produced by cAMP, although, of course, it may alter ΔG values in each state. Interestingly, ΔΔG for SthK_Cterm_ (-3.03 Kcal/mol) and SthK_Full_ (-3.10 kcal/mol) were very similar. Although the calculated energetics are model dependent, the modular gating scheme appears to capture the most salient features of SthK function and provides a context for interpreting our data.

**Figure 7.**
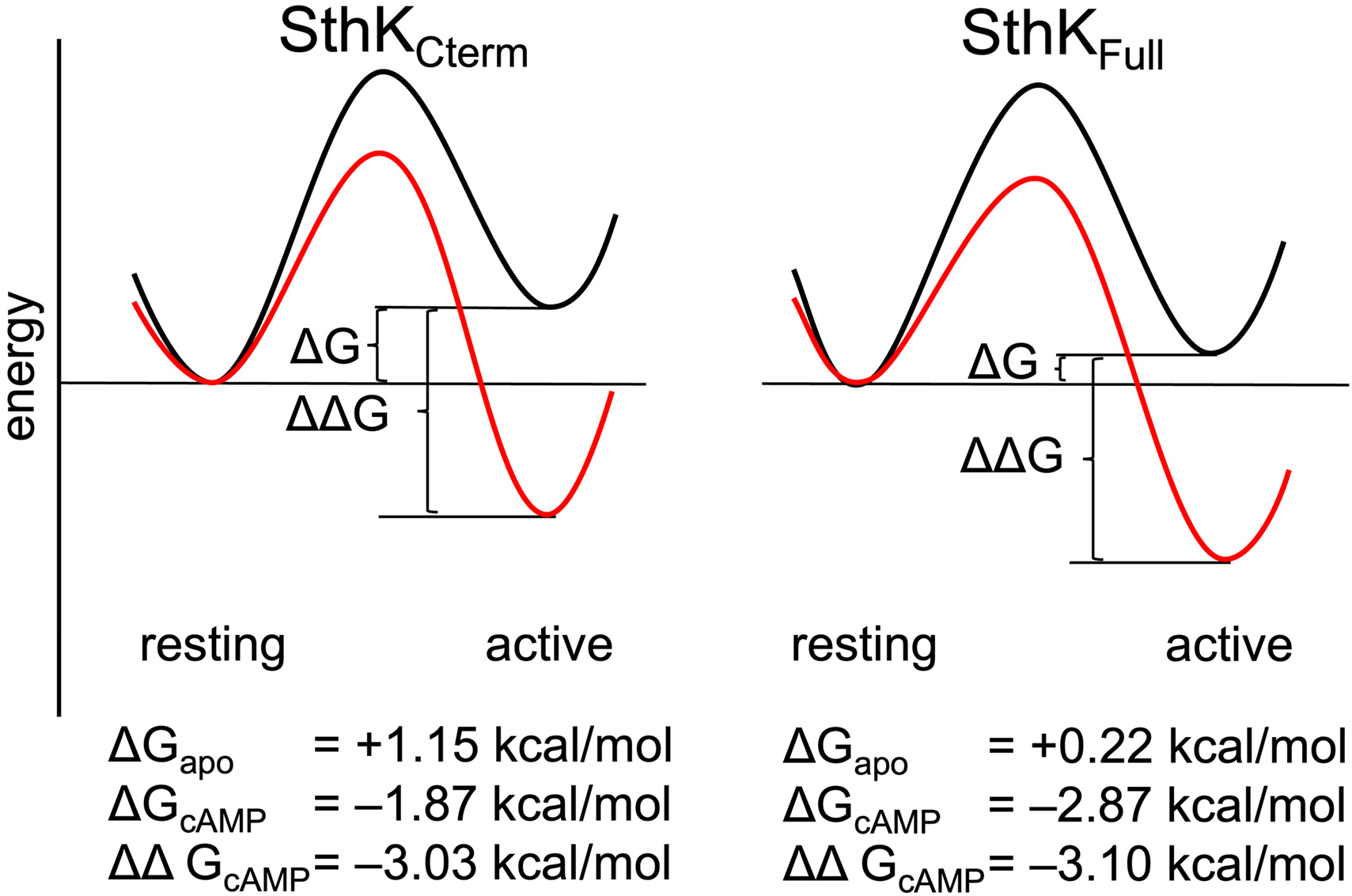
Energetics of the resting to active conformational change in the CNBD. A hypothetical reaction coordinate shows the relative energy difference between resting and active states (ΔG) and difference in free energy change (ΔΔG) without ligand (black) and with cAMP bound (red), for SthK_Cterm_ (left) and SthK_Full_ (right). Calculated ΔG and ΔΔGs values are shown in for each construct below the diagram.

## Discussion

We extended our previous work measuring conformational energetics in the SthK CNBD from a fragment containing the C-terminal C-linker and CNBD to the full-length channel complete with pore and other transmembrane domains. Here we used the same donor-acceptor pair in the CNBD as in our previous work, albeit with a different method for measuring fluorescence lifetimes. We found that the donor-acceptor distances were remarkably consistent across all constructs, within ∼1.5 Å, both in the absence of cAMP and in the presence of a saturating concentration of cAMP. The energetics of the full-length channel (with its pore and transmembrane domain), however, differed from those of the C-terminal fragment. Relative to the C-terminal fragment, the resting-to-activated conformational transition in the full-length channel was stabilized by about-1 kcal/mol, as reflected by a greater *f*_A2_ value. This stabilization of the active state relative to the resting state in the full-length channel was observed in both the absence of cAMP and in the presence of a saturating concentration of cAMP, i.e., it was not cAMP dependent. Consequently, the difference in ΔG values between the no ligand and fully-liganded conditions (ΔΔG) was the same for both the C-terminal fragment and full-length channels.

A technical challenge when working with multimeric membrane proteins is the difficulty of producing proteins with only a single labeled subunit. As a result, we measured the potential contributions of FRET between a donor on one subunit of SthK and acceptors on the other three subunits using the C-terminal fragment in which it was straightforward to mix different protein constructs. The small but significant intersubunit FRET we observed required that we consider this contribution in our FRET model with which we fit our lifetime data. After confirming that we could use an adapted FRET model for lifetime data with homotetramers of C-terminal CNBD protein in which all subunits had both donor and acceptor, we applied this method to similarly homotetrameric full-length channels. Although all fits to the data converged and gave χ^2^ values near 1, representing intersubunit donor-acceptor distances as arising from a single additional acceptor may not fully capture the intersubunit distance distribution in a tetrameric protein. Nonetheless, this approach improved our tmFRET accuracy for this donor-acceptor site and can be applied to future protein systems where FRET with multiple acceptors cannot be avoided.

Another approach would be to use a donor-acceptor pair with a shorter *R*_O_, so that the intersubunit FRET is not significant.

There are two extreme, thermodynamically equivalent models for describing the allosteric coupling between the CNBD (including the C-linker) and the pore. The active CNBD could do work on the pore to open it or the resting CNBD could do work on the pore to close it. These models, both of which are compatible with our modular gating scheme (Figure 1B), predict different effects of coupling on the system. In the first model, coupling between the pore and C-terminus makes opening of the pore more favorable and activation of the CNBD less favorable. In the second model, the coupling makes opening of the pore less favorable and activation of the CNBD more favorable. In our experiments, we assumed that deleting the pore and transmembrane domains simply eliminates the coupling of these regions to the CNBD. Our data, showing more favorable activation of the CNBD in the full-length channel containing the pore, are consistent with the second model in which the resting CNBD does work on the pore to close it. A similar conclusion was reached for HCN1 channels and spHCN channels, for which deleting the C-terminal domains gave more favorable activation of the pore in electrophysiology experiments (27–29). Our conclusions from the present study are limited because our FRET reporters were only in the CNBD and coming from a single donor-acceptor pair, with further advances requiring FRET reporters in the C-linker and pore. Additionally, we did not observe cooperativity between subunits for the CNBD conformational change, and future experiments should investigate where intersubunit cooperativity might arise in these channels. To gain a full mechanistic understanding of allostery in CNBD channels will also require measurements in native lipid bilayers and perhaps even in cells, with their complement of interacting proteins and small molecules.

## Materials and methods

### Protein expression and purification of SthK_Cterm_

The creation, expression and purification of SthK_Cterm_ protein constructs was carried out as previously described (12). Briefly, all protein constructs were expressed in B-95.ΔA *E. coli* (DE3) cells (30), lysed on an Avestin EmulsiFlex-C3 cell disruptor, and lysates were purified on Strep-Tactin high-capacity beads (IBA Biosciences) into KBT (150 mM KCl, 50 mM Tris, 10% glycerol, pH 7.9) with β-mercaptoethanol. For heterotetrameric constructs, excess of purified subunit (either SthK_Cterm_:WT or SthK_Cterm_:V416C) were added at > 7:1 ratio to the purified Acd containing constructs. For SthK_Cterm_:S361Acd-V416C homotetramers, no additional SthK_Cterm_:WT was added. All constructs were TEV cleaved to remove the MBP fusion tag and Twin-strep tag and run on ion exchange chromatography to isolate SthK_Cterm_. Protein was treated with TCEP to ensure cysteines were fully reduced, run on PD MidiTrap G-10 (Cytiva) to remove reducing reagents and exchange buffer back into KBT, and then immediately flash frozen with liquid nitrogen (LN) for storage at-80° C.

### Constructs, expression and purification of SthK_Full_

Constructs for the full-length channel SthK_Full_ were created by modifying the previously published construct of the cysteine free SthK (cfSthK) in the pCGFP vector (24). The GFP sequence fragment was removed from the C-terminal end of SthK by Gibson cloning and replaced with the sequences for a TEV protease cleavage sequence followed by a Twin-Strep-tag sequence (the same as the SthK_Cterm_ constructs). Using site-directed mutagenesis, an amber stop codon was introduced at S361 and an acceptor site was engineered by mutating V416 to cysteine to create two constructs: SthK_Full_-S361Acd and SthK_Full_-S361Acd-V416C.

These constructs were co-transformed with the AcdA9 aminoacyl tRNA synthetase/tRNA-containing plasmid (AcdA9.pDule2) (31) into B-95.ΔA *E. coli* (DE3) cells (30). 1 L cultures of transformed *E. coli* were grown in terrific broth medium at 37° C in 100 μg/ml carbenicillin and 120 μg/ml spectinomycin to an OD_600_ ∼0.4 before adding Acd (for a final concentration of 0.3 mM) and slowly lowering the temperature to 18° C (32, 33). For protein induction, 0.5 mM isopropyl β-D-1-thiogalactopyranoside (IPTG) was added, and cultures were shaken at 18° C for 20 hours followed by harvesting by centrifugation. Cell pellets were resuspended in lysis buffer (150 mM KCl, 30 mM Tris, 2 mM β-mercaptoethanol, and 10% glycerol, pH 7.9 supplemented with mini EDTA-free protease inhibitor tablets [Pierce, ThermoFisher]) and lysed on an Avestin EmulsiFlex-C3 cell disruptor 5x times at 15,000-20,000 psi. Lysate was diluted and centrifuged at 30,000x g for 30 minutes at 4° C. Clarified lysate was then spun for 1.25 hours at 200,000x g at 4° C to isolate membranes. *E. coli* membranes were then resuspended 1:2 [wt/vol] in membrane resuspension buffer (150 mM KCl, 30 mM Tris, 20% Glycerol, and 2 mM β-mercaptoethanol, pH 7.9 with an additional mini-protease inhibitor tablet) using a tissue homogenizer. Resuspended membranes were then solubilized with 1:1 [vol/vol] of solubilization buffer (150 mM KCl, 30 mM Tris, 80 mM DDM (Anatrace), 8 mM CHS (Anatrace), 2 mM β-mercaptoethanol, pH 7.9) for 1 hour and added to 1 mL of Strep-Tactin Superflow high-capacity beads (IBA Biosciences) at 4° C in a disposable column. The resin was washed with 25 mL of 2 mM β-mercaptoethanol supplemented KBT (150 mM KCl, 50 mM Tris, pH 7.9) with 1 mM DDM, 0.1 mM CHS and then 10 mL of KBT with 4 mM LMNG, 0.4 mM CHS. Detergent was allowed to exchange on the column for 30 mins at 4° C. An additional 10 mL of KBT was applied with 1 mM LMNG, 0.1 mM CHS with 1 mM tris(2-carboxyethyl)phosphine (TCEP) instead of β-mercaptoethanol. Protein was eluted from the Strep-Tactin resin with 10 mM d-Desthiobiotin (Sigma) in KBT buffer with 1mM LMNG and 0.1mM CHS and 1mM TCEP.

Detergent solubilized protein was then run on a bolt 4-12% bis-tris polyacrylamide gel (Invitrogen) for in-gel fluorescence and run analytically on a Superose 6 5/150 size exclusion chromatography (SEC) column (Cytiva) with 0.2 mM LMNG, 0.02 mM CHS to check for aggregates and protein stability. A PD-10 MidiTrap G-10 (Cytiva) desalting column was used to remove TCEP and lower detergent to 0.2 mM LMNG, 0.02 mM CHS. Protein was immediately frozen with LN and stored at-80° C for later labeling and TCSPC experiments.

### Labeling of SthK_Cterm_ and SthK_Full_ with metal acceptor for tmFRET experiments

All constructs of SthK_Cterm_ (both with cysteine and without) were incubated with 1 mM [Ru(2,2′-bpy)_2_(1,10-phenanthroline-5-maleimide)]^2+^ ([Ru(bpy)_2_phenM]^2+^, in DMSO) for 30 minutes before running on SEC to remove excess label and to isolate tetramers from monomers.

Tetrameric SthK_Cterm_ was then used directly in TCSPC experiments.

The SthK_Full_-S361Acd and SthK_Full_-S361Acd-V416C constructs were labeled with 1 mM [Ru(bpy)_2_phenM]^2+^ for 30 minutes and then cleaned up as described previously using a BioSpin6 Mini column (BioRad) (12).

### TCSPC Lifetime measurements

TCSPC lifetime data of Acd labeled protein samples (85 µL) were measured in 50 µL volume quartz cuvettes (Starna Cells, inc.). Lifetime data were measured using a PicoQuant FluoTime 300 Fluorescence Lifetime Spectrometer (PicoQuant, Berlin Germany) with 375 nm UV laser excitation and 460 nm emission, and single photon arrivals were recorded on a hybrid photomultiplier detector assembly (PMA-40). Four cuvettes were consecutively recorded, the first two with Acd labeled protein, the third with buffer-only solution equivalent to the protein sample without protein, and the fourth with diluted Ludox for measurement of the instrument response function (IRF). To each of the first three cuvettes, Adenosine 3’,5’-cyclic monophosphate sodium salt monohydrate (cAMP, Sigma Aldrich) in KBT buffer (150 mM KCl, 30 mM Tris, 10% glycerol, pH 7.9) was added at various concentrations following initial measurements without ligand. TCSPC data were acquired with the EasyTau2 measurement software (PicoQuant, Berlin Germany) and exported for analysis in Igor Pro v8 (Wavemetrics).

### FRET model of time-domain fluorescence lifetime data to obtain Gaussian distance distributions

The time-domain fluorescence lifetime data of the donor fluorophore in the presence of the acceptor were fit by model estimates for the time course of the decay, *Decay(t)*. These fits were calculated from the convolution of the measured instrument response function, *IRF(t)* of the system with model estimates of the fluorescence lifetime of the donor in the presence of the acceptor, *I_DA_(t*) and buffer-only fluorescence *I_B_(t*):

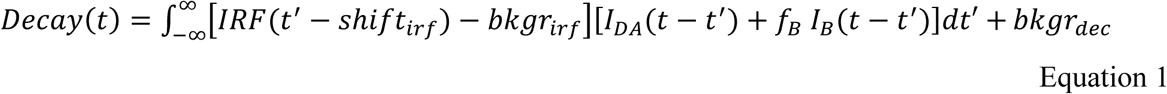

Where the variables are defined in Table I and displayed graphically in Figure 8.

**Figure 8.**
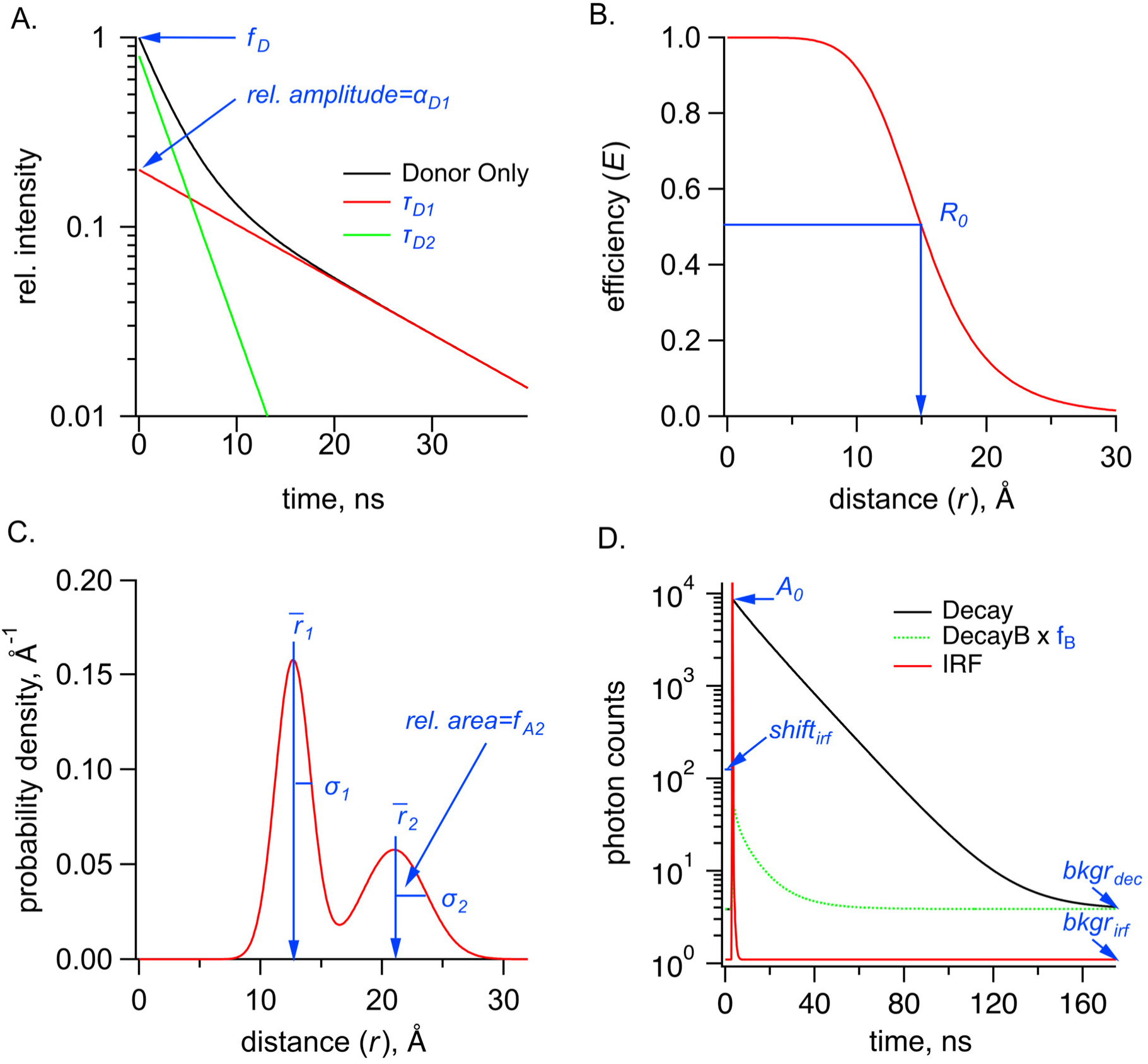
Parameters used in the FRET model, shown in blue, for TCSPC data for the sum of two Gaussians distribution. **(A)** Graph of donor-only fluorescence-lifetime decay with two exponential components with time constants (τ_D1_, τ_D2_). The amplitude fraction of the donor-only lifetime in an experiment is determined as *f*_D_. (**B)** FRET efficiency (*E*) plot as a function of distance (*r*) between donor and acceptor and the *R_0_* values for 50% FRET transfer. **(C)** Probability distribution plot of donor and acceptor distances P(*r*) showing the sum of two Gaussian distributions, each with their own average distance (*r*-_1_ and *r*-_2_), standard deviations (σ_1_ and σ_2_) and relative amplitude of the second component (*f*_A2_). **(D)** Example TCSPC data is shown with experimental measured photon count data (Decay), the instrument response function (IRF), and the buffer only decay (DecayB), with its corresponding scaling factor *f*_B_ relative to the Decay trace. Also indicated is a time-independent background of photon counts for the Decay trace, *bkgr*_dec_, as well as two parameters associated with the IRF, its background, *bkgr*_irf_, and the shift between instrument response function and measured decay, *shift*_irf_. Parameter *A_O_* is the amplitude of the FRET model estimate of the fluorescence lifetime of the donor (in photon counts).

**Table I.**

| Variable | Description |
| --- | --- |
| $t$ | Time of photon emission (in s) |
| $r$ | Distance between the donor and acceptor (in Å) |
| $Decay(t)$ | Time-dependent decay of the donor fluorescence (in counts) |
| $DecayB(t)$ | Time-dependent decay of the buffer-only fluorescence (in counts) |
| $I_{DA}(t)$ | Model estimate of the fluorescence lifetime of the donor in the presence of the acceptor (in counts) |
| $I_B(t)$ | Model estimate of the fluorescence lifetime of a buffer-only sample (in counts) |
| $IRF(t)$ | Measured instrument response function (in counts) |
| $bkg_{dec}$ | Time-independent decay background (in counts) |
| $bkg_{irf}$ | Time-independent background of the instrument response function (in counts) |
| $shift_{irf}$ | Time shift between the instrument response function and the measured decay (in ps) |
| $f_B$ | Scaling of the buffer fluorescence |
| $A_0$ | Amplitude of the model estimate of the fluorescence lifetime of the donor (in counts) |
| $f_D$ | Fraction of donor-only fluorescence in the sample |
| $\alpha_{Di}$ | Fraction of the $i$th component of the donor-only fluorescence (in counts) |
| $\tau_{Di}$ | Time constant of the $i$th component of the donor-only fluorescence (in s) |
| $\alpha_{Bi}$ | Amplitude of the $i$th component of the buffer fluorescence (in counts) |
| $\tau_{Bi}$ | Time constant of the $i$th component of the buffer fluorescence (in s) |
| $R_0$ | Characteristic distance between donor and acceptor producing 50% FRET efficiency |
| $\rho(r)$ | Probability distance distribution of the donor and acceptor distances |
| $f_{Ai}$ | Fraction of the $i$th component of the probability distance distribution |
| $\bar{r}_i$ | Average distance of the $i$th component of the donor-only fluorescence (in Å) |
| $\sigma_i$ | Standard deviation of the $i$th component of the donor-only fluorescence (in Å) |

The model for fluorescence lifetime assumes a donor-only fluorescence lifetime with one or two exponential components, and FRET between a donor and acceptor separated by one or two Gaussian-distributed distances, similar to that previously described for frequency domain measurements (9, 18, 21, 34). This model predicts the following relationship for the fluorescence lifetime of the donor in the presence of acceptor, *I_DA_(t*):

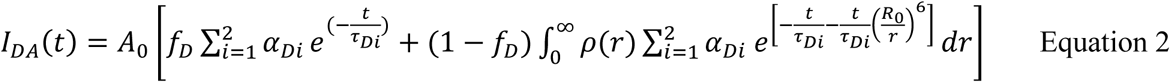

Where α_D1_ + α_D2_ = 1.

The density distribution of donor-acceptor distances, ρ(*r*), was assumed to be the sum of up to two Gaussians:

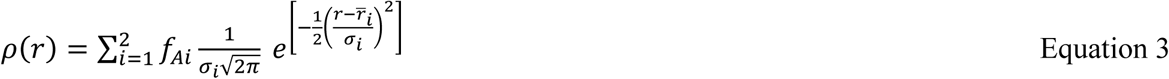

where *f*_A1_ + *f*_A2_ = 1.

The decay time course of the buffer-only sample, *DecayB(t*), was fit with a convolution of the measured instrument response function, *IRF(t)* with a multi-exponential model for the fluorescence lifetime, *I_B_(t*):

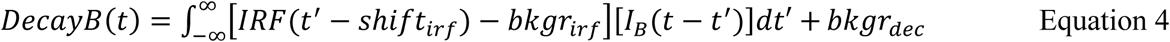

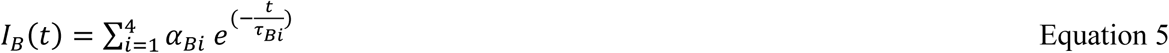

This model for the buffer-only fluorescence lifetime was then added to the model for the sample lifetime before convolving with the *IRF(t)* (Equation 1).

This FRET model for fluorescence lifetimes and FRET was implemented in Igor Pro v8 (Wavemetrics, Lake Oswego, OR) (code available at https://github.com/zagotta/TDlifetime_program). The FRET model was either fit, with χ^2^minimization, assuming a standard deviation equal to 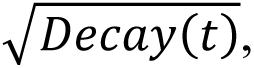 to individual decay time courses, or globally fit to multiple decays with the sample under different conditions as indicated in the text. There are 15 parameters in the parameter vector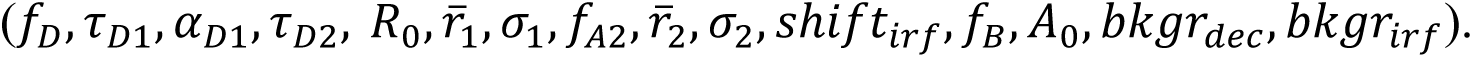 was fixed to a value previously determined from emission spectra of the donor and absorbance spectra of the acceptor using the Förster equation assuming *k*^2^ = 2/3. *f*_A2_,was generally fixed to 0 or 1 for fits assuming a single Gaussian distance distribution but was allowed to vary when multiple distance components were expected (e.g. with subsaturating ligand concentrations). *f*_B_ was fixed to the acquisition time for the experimental sample divided by the acquisition time for the buffer-only sample. And *bkgr*_irf_was always fixed to 0 as the average background photon count in the *IRF(t)* was <<1.

For the individual decay fits to the donor-only sample, *f*_D_ was set to 1, and only 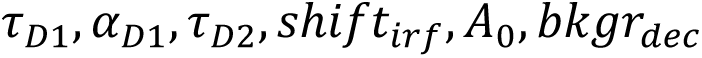 were allowed to vary. For the individual decay fits to the donor+acceptor samples, 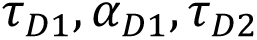 were constrained to the previously determined values from the donor-only sample, and 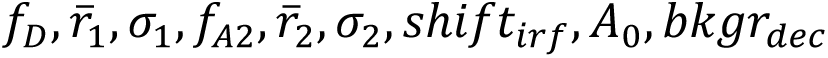 were varied. For global fits to multiple decays with the sample under different conditions, multiple parameters were constrained to be the same across some or all of the decay fits, as indicated in the text.

To assess the resolvability of our parameters in our TCSPC FRET model, we generated χ^2^plots for the key parameters (Figure 8-Supplemental Figure 1). All parameters we tested reliably converged on a minimized reduced χ^2^ value (∼1), although the σ parameters produced plots with much shallower slopes and were noticeably less well determined than the other parameters.

Additionally, for the parameters expected to have highest correlation, we calculated reduced χ^2^surfaces (Figure 8-supplemental figure 2). A small correlation was observed between σ_1_ and the fraction donor-only, *f*_D_, and between the average distance 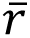_1_ and *f*_D_. As a result, we were careful about our interpretation of *σ* values. Overall, the close agreement of 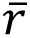, σ, and *f*_A2_ values obtained from independent measurements and between TCSPC and frequency domain measurements indicates that time-resolved tmFRET is robust across different methods of lifetime measurements.

### Adaptation of the FRET model of time-domain fluorescence lifetime data to include inter-subunit FRET

The relationship for the normalized fluorescence lifetime of the donor when there is a single acceptor is given by a simplified version of Equation 2:

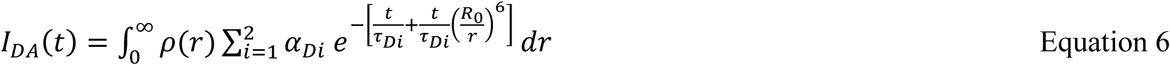

Adding a second FRET acceptor with the same *R_O_* and a new distance *r*_2_ gives the following equation for the normalized fluorescence lifetime:

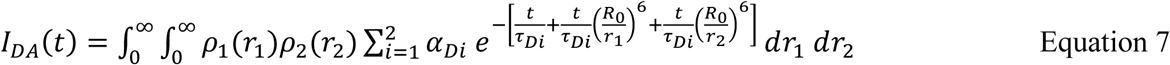

where ρ_1_(*r*_1_) is the distance distribution of the first acceptor and ρ_2_(*r*_2_) is the distance distribution of the second FRET acceptor. Since the distance distribution of each acceptor is independent of the distance to the other acceptor, this equation can be rewritten as the product of integrals as follows:

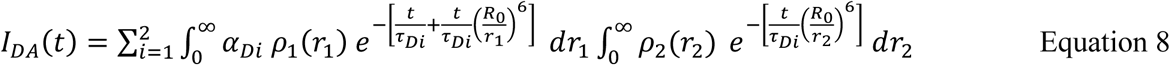

To determine the fluorescence lifetime with both an intrasubunit acceptor and an intersubunit acceptor, the experimentally measured intersubunit FRET (Figure 3) was first fit with the lifetime model from Equation 2 assuming a single Gaussian distribution for the resting and active states. For simplicity, the FRET distance distribution from three potential intersubunit acceptors was approximated by one Gaussian which largely reflects the closest of the three acceptors. In addition, the fractions of the Gaussian for the activated state (*f*_A2_) were fixed at values of 0.2 and 0.8 for the apo and cAMP conditions, respectively. The obtained 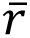_1_, σ_1_, 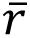_2_, σ_2_ values from this fit were then used as fixed parameters for the distribution ρ_2_(*r*_2_) in Equation 8 when fitting homotetrameric SthK_Cterm_ and SthK_Full_ lifetime data. While the ρ_2_(*r*_2_) *r̅s* and *σs* of the intersubunit distance distribution were held constant, the ratio between these Gaussians (*f*_A2_) was set to the same *f*_A2_ value as that of ρ_1_(*r*_1_). In this way, 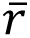_1_, σ_1_, 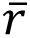_2_, σ_2_, *and f*_A2_were determined for the intrasubunit distance distribution ρ_1_(*r*_1_).

### Free Energy and Dose-Response Calculations

Free energies were calculated using the following equations:

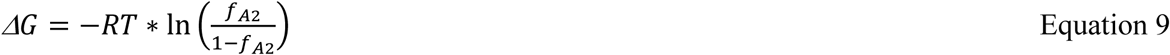

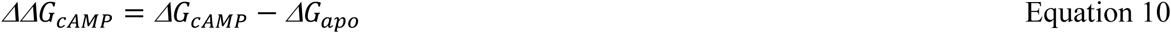

where *R* is the universal gas constant, *T* is the absolute temperature (K), and *f*_A2_ and 1 - *f*_A2_ are the proportion of molecules in the active and resting states respectively.

Dose-response relations (Figure 2F and Figure 6F) were fit by the following quadratic equation:

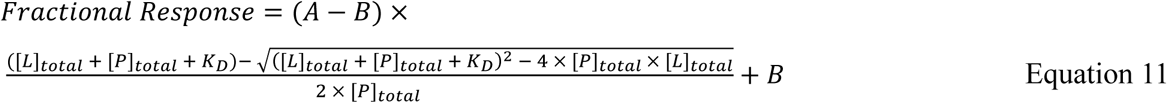

Where *K_D_* is the dissociation constant for cAMP, *[L]_total_* is the total ligand (cAMP) concentration, [*P*]_total_ is the total protein concentration, *A* is the *f*_A2_ value at saturating ligand concentrations and *B* is the *f*_A2_ value without ligand. The above parameters were obtained from least squares fits to the data, and the protein concentration [*P*]_total_ was corroborated via A_280_ and Bradford assay for each sample.

### chiLife Predictions

Computational predictions of the possible rotameric positions for the donor and acceptor labels were made with chiLife using the accessible-volume sampling method as previously described (19, 23). Acd and [Ru(bpy)_2_phenM]^2+^ were added as custom labels in chiLife and modeled onto the cryo-EM structure of the full-length SthK closed state (25) (PDB: 7RSH, residues 225 to 416) and the X-ray crystallography structure of the cAMP bound SthK C-terminal fragment (26) (PDB: 4D7T). For each donor-acceptor pair, labels were superimposed at indicated residue positions and 10,000 possible rotamers were modeled. Rotamers resulting in internal clashes (<2 Å) were removed and external clashes evaluated as previously described (23). Donor-acceptor distance distributions were calculated between the remaining (∼500-2000) label rotamers for each donor-acceptor pair. Inter-subunit distances were calculated as distances between one Acd molecule and the modeled acceptor rotamers on each of the other three subunits. Structural representations were made using Pymol (V3.0 Schrödinger, LLC).

## Acknowledgements

We thank the Oregon State University GCE4ALL (Center for Genetic Code Expansion for All) for their long-standing collaboration, and Drs. James Petersson, and Kyle D Shaffer (University of Pennsylvania) for excellent technical support with Acd. Thanks to Drs. Max Tessmer and Stefan Stoll (University of Washington) with technical assistance with chiLife. We also thank all members of the S.G. and W.N.Z. laboratories for helpful conversations and support. Research reported in this publication was supported by the National Institutes of Health under award numbers R35GM145225 (to S.E.G.), R35GM148137 and R01EY010329 (to W.N.Z.), T32GM008268 and T32EY007031 (to P.E.).

## Data Sharing Plan

Source data files have been included with the final manuscript. Code used for data analysis has been posted to https://github.com/zagotta/TDlifetime_program.

## Supplemental Figures

**Figure 8 – supplemental figure 1.**
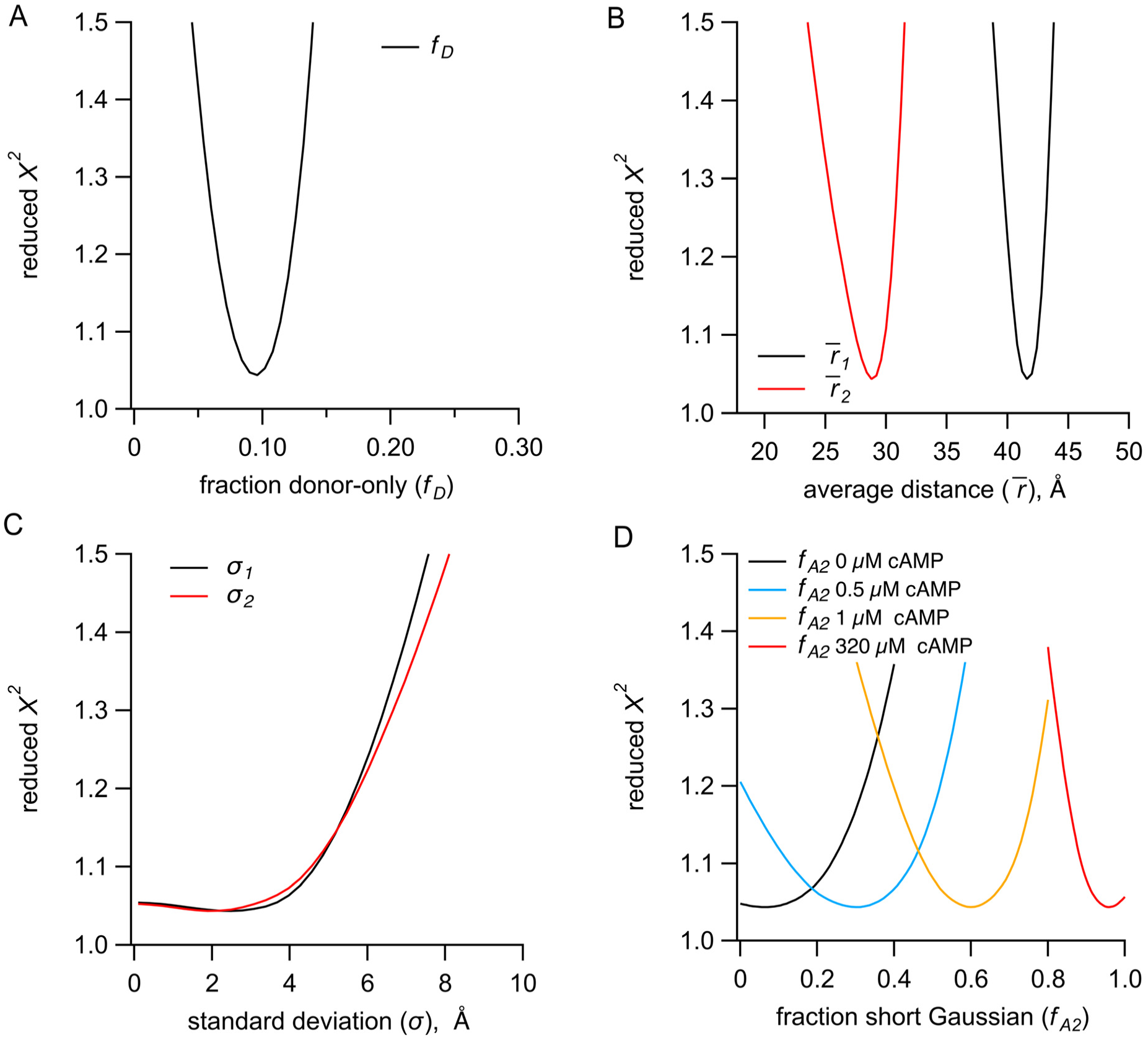
A) Identifiability of parameters in the sum of two Gaussian distributions model for representative TCSPC data. **(A)** Dependence of the reduced χ^2^ values on the fraction of donor-only component (*f*_D_). **(B-C)** Dependence of the reduced χ^2^ values on the Gaussian average distances (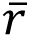) (B) and standard deviations (σ) (C) for the resting (black) and active states (red). **(D)** Dependence of the reduced χ^2^ values on the parameter *f*_A2_for the conditions of apo, 0.5 μM, 1 μM and 320 μM cAMP.

**Figure 8 – supplemental figure 2.**
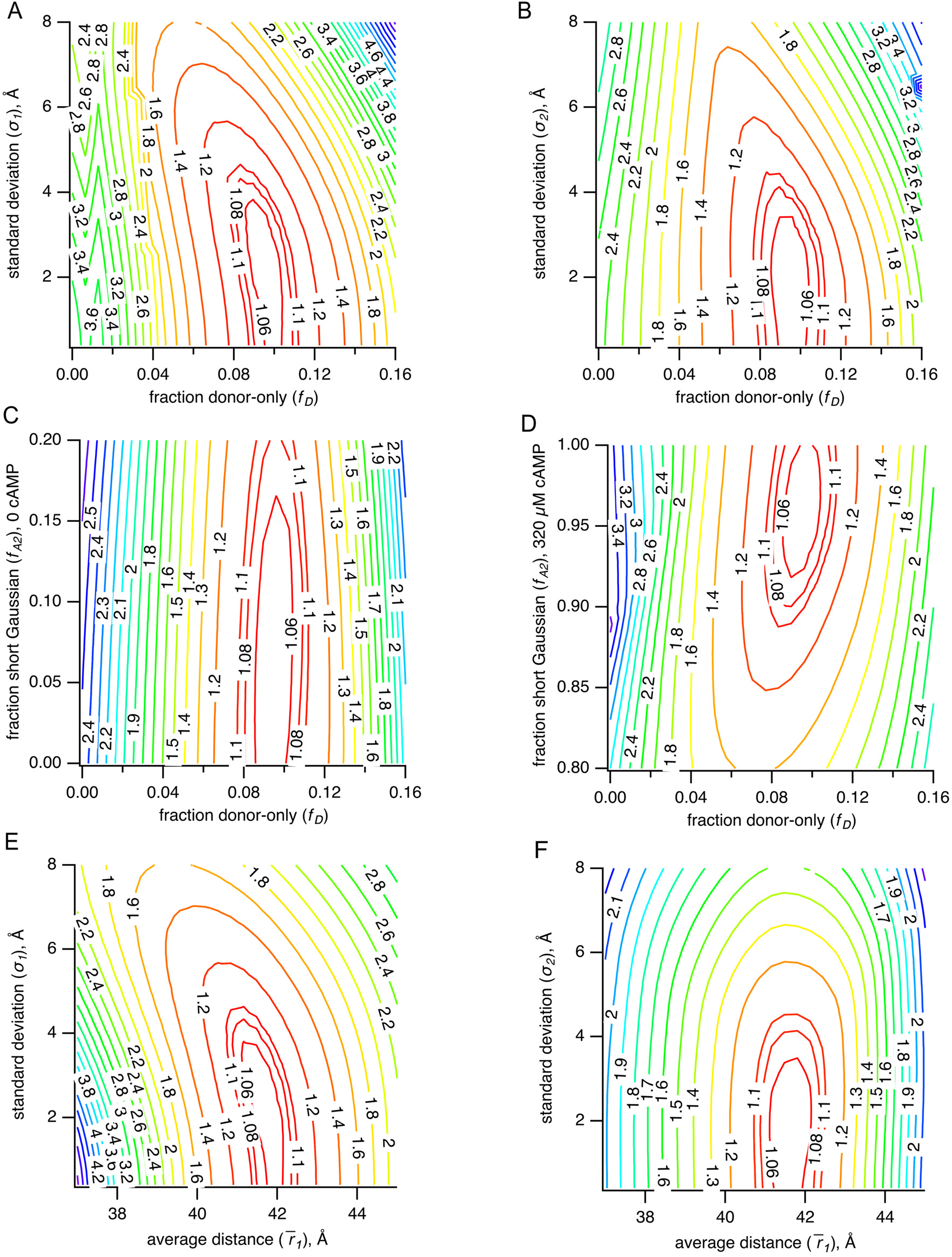
(A-F) Parameter correlations for representative TCSPC data of χ^2^ values in 3 dimensions. Plots of χ^2^ values vs. σ_1_ and *f*_D_ (A), vs. σ_2_ and *f*_D_ (B), vs. *f*_A2_ in 0 μM cAMP and *f*_D_ (C), vs. *f*_A2_ in 320 μM cAMP and *f*_D_ (D), vs σ_1_ and 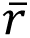_1_ (E) and vs. σ_2_ and 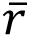_1_(F).

## References

1. K. B. Craven, W. N. Zagotta, CNG and HCN channels: Two peas, one pod. Annu Rev Physiol 68, 375–401 (2006).

2. C. He, F. Chen, B. Li, Z. Hu, Neurophysiology of HCN channels: From cellular functions to multiple regulations. Prog Neurobiol 112, 1–23 (2014).

3. K. Matulef, W. N. Zagotta, Cyclic Nucleotide-Gated Ion Channels. Annu Rev Cell Dev Biol 19, 23–44 (2003).

4. E. G. B. Evans, J. L. W. Morgan, F. DiMaio, W. N. Zagotta, S. Stoll, Allosteric conformational change of a cyclic nucleotide-gated ion channel revealed by DEER spectroscopy. Proc Natl Acad Sci U S A 117, 10839–10847 (2020).

5. Z. M. James, W. N. Zagotta, Structural insights into the mechanisms of CNBD channel function. Journal of General Physiology 150, 225–244 (2018).

6. K. B. Craven, N. B. Olivier, W. N. Zagotta, C-terminal movement during gating in cyclic nucleotide-modulated channels. J Biol Chem 283, 14728–14738 (2008).

7. M. Brams, J. Kusch, R. Spurny, K. Benndorf, C. Ulens, Family of prokaryote cyclic nucleotide-modulated ion channels. Proc Natl Acad Sci U S A 111, 7855–7860 (2014).

8. P. A. M. Schmidpeter, C. M. Nimigean, Correlating ion channel structure and function, 1st Ed. (Elsevier Inc., 2021).

9. J. Rheinberger, X. Gao, P. A. M. Schmidpeter, C. M. Nimigean, Ligand discrimination and gating in cyclic nucleotide-gated ion channels from apo and partial agonist-bound cryo-EM structures. Elife 7, 1–25 (2018).

10. P. A. M. Schmidpeter, X. Gao, V. Uphadyay, J. Rheinberger, C. M. Nimigean, Ligand binding and activation properties of the purified bacterial cyclic nucleotide-gated channel SthK. Journal of General Physiology 150, 821–834 (2018).

11. A. Marchesi, et al., An iris diaphragm mechanism to gate a cyclic nucleotide-gated ion channel. Nat Commun 9 (2018).

12. P. Eggan, S. E. Gordon, W. N. Zagotta, Ligand-Coupled Conformational Changes in a Cyclic Nucleotide-Gated Ion Channel Revealed by Time-Resolved Transition Metal Ion FRET. Elife 2024.04.25.591185 (2024). 10.1101/2024.04.25.591185.

13. F. T. Horrigan, R. W. Aldrich, Coupling between voltage sensor activation, Ca2+ binding and channel opening in large conductance (BK) potassium channels. J Gen Physiol 120, 267–305 (2002).

14. H. N. Motlagh, J. O. Wrabl, J. Li, V. J. Hilser, The ensemble nature of allostery. Nature 508, 331–339 (2014).

15. V. J. Hilser, J. O. Wrabl, H. N. Motlagh, Structural and energetic basis of allostery. Annu Rev Biophys 41, 585–609 (2012).

16. K. B. Craven, W. N. Zagotta, Salt bridges and gating in the COOH-terminal region of HCN2 and CNGA1 channels. J Gen Physiol 124, 663–677 (2004).

17. H. A. DeBerg, P. S. Brzovic, G. E. Flynn, W. N. Zagotta, S. Stoll, Structure and energetics of allosteric regulation of HCN2 ion channels by cyclic nucleotides. Journal of Biological Chemistry 291, 371–381 (2016).

18. E. Haas, M. Wilchek, E. Katchalski-Katzir, I. Z. Steinberg, Distribution of end-to-end distances of oligopeptides in solution as estimated by energy transfer. Proc Natl Acad Sci U S A 72, 1807–1811 (1975).

19. W. N. Zagotta, et al., Measuring conformational equilibria in allosteric proteins with time-resolved tmFRET. Biophys J 2023.10.09.561594 (2024). 10.1016/j.bpj.2024.01.033.

20. W. N. Zagotta, et al., An improved fluorescent noncanonical amino acid for measuring conformational distributions using time-resolved transition metal ion FRET. Elife 10, 2021.05.10.443484 (2021).

21. J. R. Lakowicz, Principles of fluorescence spectroscopy (2006).

22. S. E. Gordon, et al., Long-distance tmFRET using bipyridyl-and phenanthroline-based ligands. Biophys J 2023.10.09.561591 (2024). 10.1016/j.bpj.2024.01.034.

23. M. H. Tessmer, S. Stoll, chiLife: An open-source Python package for in silico spin labeling and integrative protein modeling. PLoS Comput Biol 19, 1–16 (2023).

24. J. L. W. Morgan, E. G. B. Evans, W. N. Zagotta, Functional characterization and optimization of a bacterial cyclic nucleotide-gated channel. Journal of Biological Chemistry 294, 7503–7515 (2019).

25. X. Gao, et al., Gating intermediates reveal inhibitory role of the voltage sensor in a cyclic nucleotide-modulated ion channel. Nat Commun 13 (2022).

26. D. Kesters, et al., Structure of the SthK carboxy-terminal region reveals a gating mechanism for cyclic nucleotide-modulated ion channels. PLoS One 10, 1–12 (2015).

27. S. Vemana, S. Pandey, H. P. Larsson, S4 movement in a mammalian HCN channel. J Gen Physiol 123, 21–32 (2004).

28. G. E. Flynn, W. N. Zagotta, Molecular mechanism underlying phosphatidylinositol 4,5-bisphosphate-induced inhibition of SpIH channels. Journal of Biological Chemistry 286, 15535–15542 (2011).

29. B. J. Wainger, M. DeGennaro, B. Santoro, S. A. Siegelbaum, G. R. Tibbs, Molecular mechanism of cAMP modulation of HCN pacemaker channels. Nature 411, 805–810 (2001).

30. T. Mukai, et al., Highly reproductive Escherichia coli cells with no specific assignment to the UAG codon. Sci Rep 5, 1–9 (2015).

31. I. Sungwienwong, et al., Improving target amino acid selectivity in a permissive aminoacyl tRNA synthetase through counter-selection. Org Biomol Chem 15, 3603–3610 (2017).

32. L. C. Speight, et al., Efficient synthesis and in vivo incorporation of acridon-2-ylalanine, a fluorescent amino acid for lifetime and Förster resonance energy transfer/luminescence resonance energy transfer studies. J Am Chem Soc 135, 18806–18814 (2013).

33. C. M. Jones, Y. Venkatesh, E. J. Petersson, Protein labeling for FRET with methoxycoumarin and acridonylalanine, 1st Ed. (Elsevier Inc., 2020).

34. A. Grinvald, E. Haas, I. Z. Steinberg, Evaluation of the Distribution of Distances Between Energy Donors and Acceptors by Fluorescence Decay. Proceedings of the National Academy of Sciences 69, 2273–2277 (1972).

